# River phosphorus cycling constrains lake cyanobacteria blooms

**DOI:** 10.1101/2021.05.03.442426

**Authors:** Whitney M. King, Susan E. Curless, James M. Hood

## Abstract

Bioavailable phosphorus exports from rivers during high flow often fuel downstream harmful cyanobacterial blooms; yet whether river phosphorus cycles affect these exports is unclear. Here, we examined river phosphorus cycling during high flow events in a large agricultural watershed that drives cyanobacterial blooms in Lake Erie. We show that between 2003 and 2019 river phosphorus cycles, through phosphorus sorption, reduced bioavailable phosphorus exports by 24%, potentially constraining Lake Erie cyanobacterial blooms by 61%. Over the last 45-years, phosphorus sorption has declined with suspended sediment exports due to increases in soil-erosion-minimizing agricultural practices, likely contributing to recent cyanobacterial blooms. In this, and likely other agricultural watersheds, rivers perform an unrecognized ecosystem service during high flow creating field-river-lake linkages that need to be incorporated into phosphorus management.

## Main Text

Cyanobacterial harmful blooms (cyanoHABs), which have been increasing worldwide, negatively impact freshwater ecosystems, human livelihoods, and human health (*1, 2*). One primary cause of cyanoHABs are phosphorus (P) losses from agricultural production, particularly the loss of dissolved reactive P (DRP) which is far more bioavailable to cyanobacteria than sediment P (*3*). Thus, it is a global environmental priority to understand and reduce watershed P exports (*4*). Yet, P management is complicated by uncertainty associated with P retention and transformation during downstream transport through river networks (*5, 6*). In streams and rivers, biogeochemical processes alter the magnitude and bioavailability (i.e., DRP versus sediment P) of P transported downstream exerting controls on P exports to recipient water bodies during low flows (*7*). Yet, P and suspended sediment concentrations increase with discharge and most P and sediment export occurs at high flows (*8–10*). Yet, we know little about P cycling during high flows (*7, 11*); constraining our ability to successfully manage P pollution and cyanoHABs.

One potentially important aspect of river P cycling during high flows is the abiotic process of P sorption-desorption by suspended sediment particles, which can increase or decrease DRP concentrations and, therefore, the bioavailability of P exports to cyanoHABs. P sorption-desorption during high flows, which is rarely quantified, is perceived to be unimportant at the watershed scale because the rapid downstream transport limits the time for sorption-desorption to influence DRP exports (Fig. 1a) (*7, 12*). Yet, in many river systems rainfall driven surface water runoff travels hours to days from catchment headwater streams to recipient water bodies (*13, 14*), allowing sufficient time for P sorption-desorption to influence DRP exports (*15*). Unfortunately, it is unknown how much P sorption-desorption affects DRP exports during high flows and, thus, subsequently mediates cyanoHABs in recipient ecosystems.

**Figure 1.**
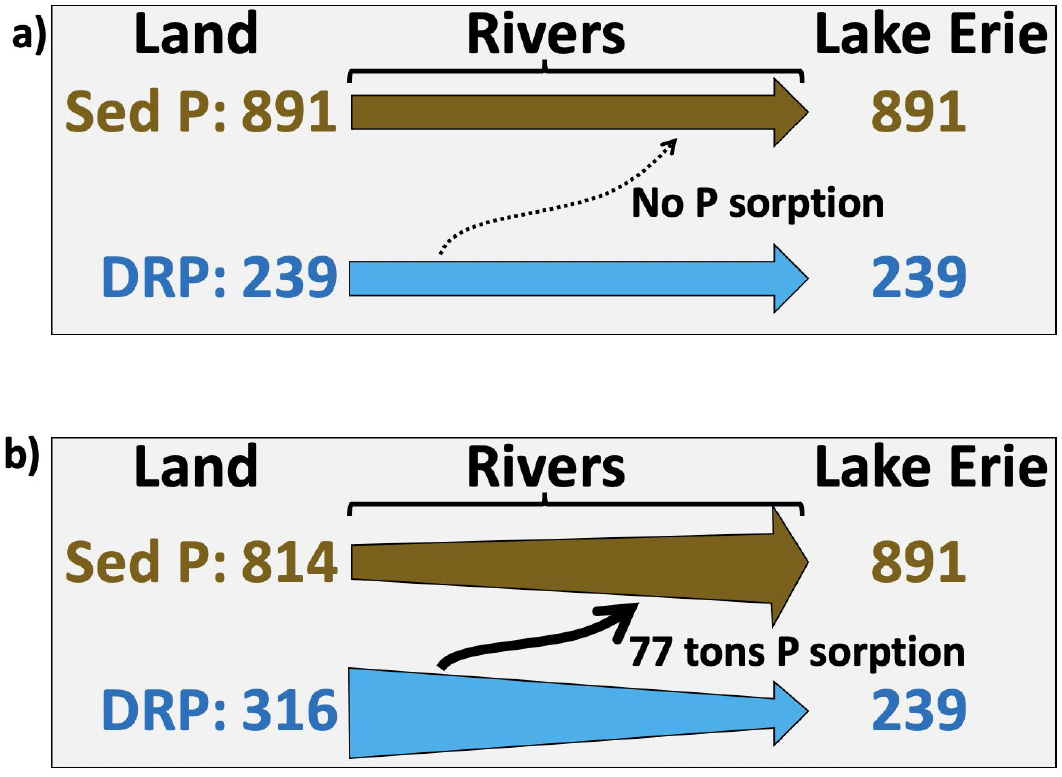
Present (a) and new (b) conceptual model of the role of streams and rivers in shaping P exports during high flow. The present model conceptualizes these systems as pipes transporting DRP and sediment P downstream from land to recipient waters. Our findings recognize the role of P sorption in changing the form of P exports from DRP to sediment P, and thus decreasing P bioavailability. Values are means of March-June high flow event loads or P sorption (metric tons P) between 2003 and 2019.

Our poor understanding of P cycling during high flows may mask an important watershed-scale process that could mediate the influence of agricultural practices on cyanoHABs, complicating P management (*5, 6*). During high flows, the magnitude of P sorption-desorption is likely determined by DRP concentrations, suspended sediment concentration, and river discharge (*16, 17*). These factors control the magnitude of P sorption-desorption in a volume of river water and ultimately, in conjunction with hydrologic travel time, the total net P sorption-desorption within a river network. Thus, controls on these factors (e.g., climate or agricultural management practices) impact the magnitude and direction of P sorption-desorption (*18, 19*) and the bioavailability of exports to recipient ecosystems.

Here, we examined the influence of P sorption-desorption during high flows on DRP exports and cyanoHABs. We hypothesized that during high flow, suspended sediment concentration shapes the magnitude of P sorption-desorption at the watershed scale altering the bioavailability of P exports and ultimately cyanoHABs severity in recipient ecosystems. We evaluated this hypothesis by measuring the magnitude and controls of P sorption-desorption by suspended sediment during high flow events. Then, we scaled up those processes to the tributary and river network level to estimate the impact of P sorption-desorption on cyanoHABs in Lake Erie.

### P sorption in tributaries

We tested our hypothesis in the agricultural-dominated Maumee River watershed which drains into Lake Erie. DRP exports between March and June from the Maumee watershed are primarily responsible for the severity of reoccurring Lake Erie cyanoHABs (*3, 20, 21*). We focus on a March–June window, as opposed to the March–July window used in many cyanoHABs forecast models (*22, 23*), because it overlaps with our high flow sampling and represents the majority of March–July DRP loads (Fig. S1). On average, 77% of Maumee River March–June DRP exports occur during high flows (<25% flow exceedance). Thus, P sorption-desorption by suspended sediment during high flow events could shape DRP exports to Lake Erie and cyanoHABs.

To better understand the influence of P cycling on DRP exports during high flow events, we measured three parameters that combine to determine the ecosystem level impact of P sorption-desorption: P sorption-desorption, equilibrium P concentration (ECP_0_), and P sorption capacity. EPC_0_ is the DRP concentration separating P sorption (DRP > EPC_0_) and desorption (DRP < EPC_0_) and predicts whether suspended sediment are a P source (desorption) or sink (sorption). P sorption capacity reflects the total P mass suspended sediment particles can sorb. In six tributaries of the Maumee River (Ohio, USA; Fig. S2; Table S1), we sampled 13 storm events between January and June 2019 (Fig. S3). Eighty-six percent of our samples were collected during high flows (i.e., Fig. S4).

Suspended sediment sorbed P in 98% of our measurements (77 of 78 samples; DRP > EPC_0_), although P sorption rates varied from 0.001 to 72.5 mg P m^−3^ h^−1^. Sorption rates varied among streams and increased with suspended sediment concentrations (R^2^ = 0.81, p-value < 0.001; Fig. 2; Table S2) and discharge (R^2^ = 0.69, p-value < 0.001), which were strongly correlated (r = 0.64).

**Figure 2.**
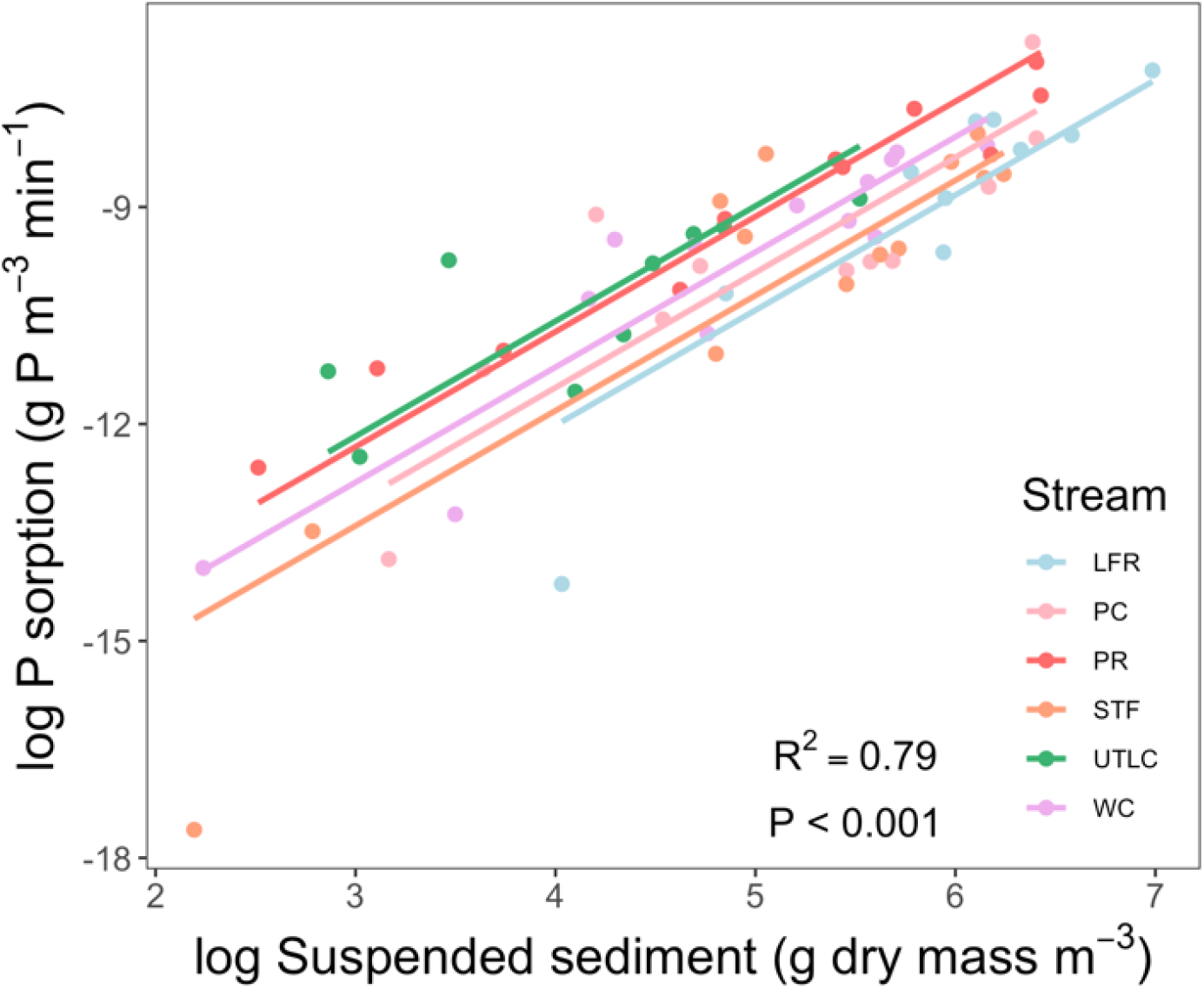
Phosphorus sorption increased with suspended sediment concentrations in six Maumee River tributaries.

Our results indicate that the 40% reductions in spring DRP load required by a binational P management agreement (*22*), would likely not alter DRP source-sink dynamics in these tributaries during high flow. Tributary DRP concentrations during high flow averaged ~800% higher than EPC_0_ indicating that a 40% reduction in DRP concentration would likely not lead to P desorption by suspended sediment. Furthermore, EPC_0_ increased with DRP concentrations with a positive intercept and slope (R^2^ = 0.48, p-value < 0.001; Fig. S5) indicating that declines in DRP concentrations might lower EPC_0_ but not alter P source-sink dynamics.

### Tributary P sorption decreases DRP exports

To estimate P sorption in tributaries, we combined P sorption parameters with discharge, DRP, and suspended sediment data from three tributaries with sufficient DRP and suspended sediment data. We focused on high flow events between March and June 2019. We estimated the magnitude of P sorption by suspended sediment exported from these tributaries in four steps. First, we calculated the potential mass-specific P sorption as the P sorbed per gram suspended sediment during transport between the tributaries and the furthest downstream Maumee River monitoring station (Waterville, OH; 26 km from Lake Erie; Fig. S2). Next, we constrained P sorption to not exceed the maximum capacity of suspended sediment to bind P (P sorption capacity). Then, we calculated total P sorption as the product of mass-specific sorption rates and suspended sediment loads. We constrained total P sorption so that it could not decrease DRP concentrations below the threshold at which sediment switch from a P sink to a source (i.e., EPC_0_). Finally, we integrated total P sorption across March–June high flow events during 2019.

Tributary P sorption substantially reduced DRP exports resulting in a significant and, until now, unrecognized ecosystem service. Suspended sediment in these tributaries sorbed 31 – 45% of observed DRP exports (0.1–0.2 tons P; Fig. 3a).

**Figure 3.**
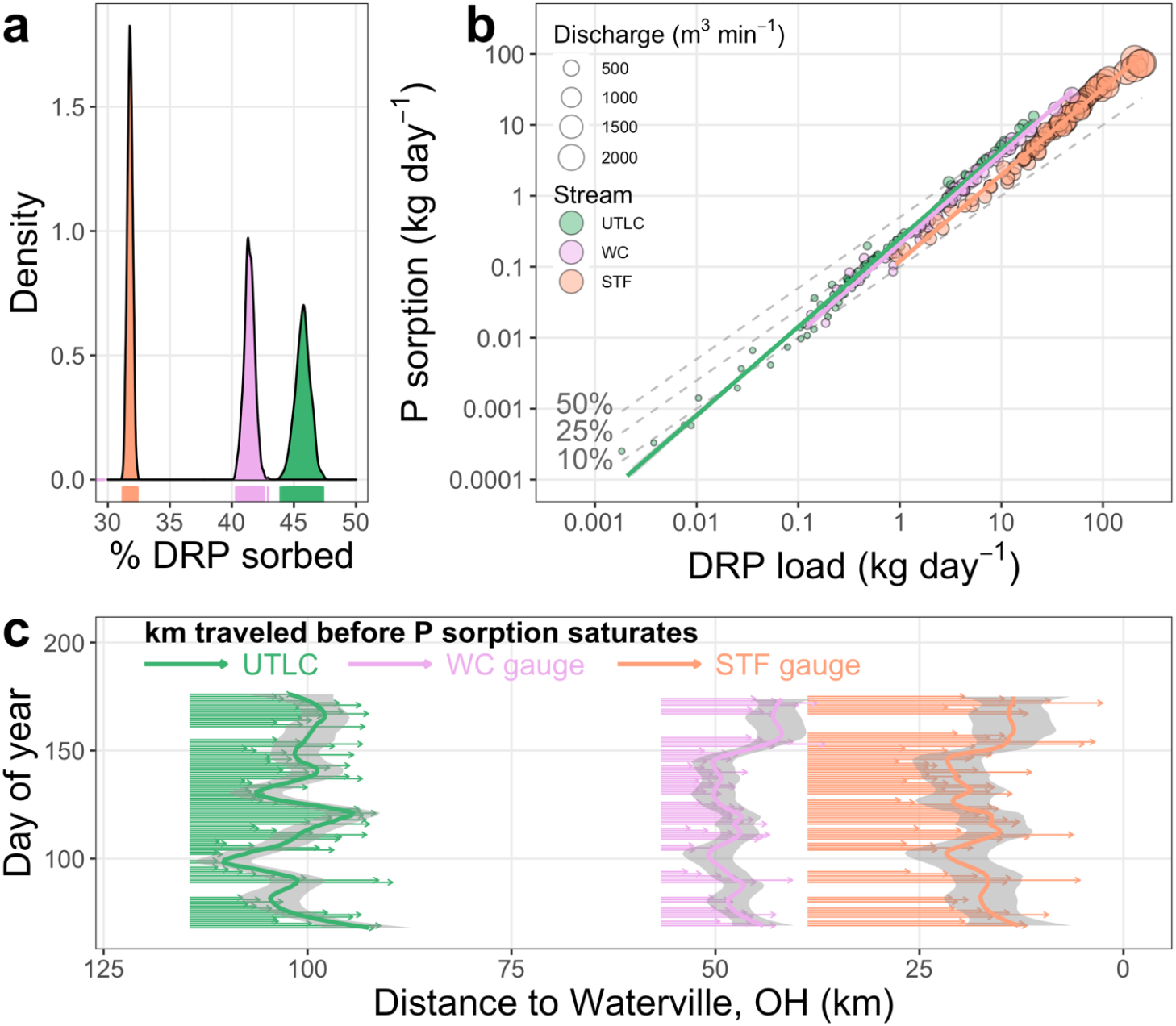
Characteristics of P sorption in three Maumee River tributaries. **a**, The percent of DRP sorbed between March and June 2019 at high flows (< 25% flow exceedance). **b**, Phosphorus sorption increases more rapidly than DRP loading, indicating that the percent of DRP sorbed (dashed grey lines) increases from <10% to >50% with both DRP loads, discharge (point size), and suspended sediment loads (not shown). **c**, The daily mean distance suspended sediment travel during high flow before P sorption saturates (each arrow) is shorter than the distance to Lake Erie (Waterville, OH is 26 km from Lake Erie). Splines are a LOESS fit to aid visualization.

Daily tributary P sorption increased with discharge and DRP loads (Fig. 3b) as well as suspended sediment loads (not shown); three tightly linked exports from agricultural watersheds. As a result, the daily percent of DRP exports sorbed increased from ~10% to 50% with discharge (Fig. 3b). Sediment P sorption saturated before the sediment had traveled an average of 14.7 km, 11 to 55% of the distance to the downstream Maumee River monitoring station (Fig. 3c). Thus, P sorption reactions reached equilibrium well before sediment were exported from the Maumee watershed, indicating that suspended sediment originating from most of the Maumee watershed have ample time during transport to shape DRP exports.

### P sorption constrains Lake Erie cyanoHABs

To investigate the effect of P sorption within the Maumee River network on Lake Erie DRP loads and cyanoHABs, we combined our sorption measurements with 1975 to 2019 discharge and water quality data from the Maumee River monitoring station at Waterville, OH (Fig. S2; Table S1). We estimated P sorption within the Maumee River system by combining data on Maumee River suspended sediment loads and watershed-scale mass-specific sorption rates from the tributaries. Our assumption that P sorption by suspended sediment did not change substantially between 1975 and 2019 is supported by comparison of recent (Fig. 2) (Fig. 2; *24*) and older (i.e., 1978 (*25, 26*)) research indicating that suspended sediment is enriched in P relative to agricultural soils, sorbed P during high flows, and had similar P bioavailability (*3*). Here, we integrate all estimates for each year across March–June high flow events; all subsequent measurements focus on this window.

During March–June high flow events, suspended sediment sorbed 17.4 – 164.8 tons P per year or 20–342% of observed DRP loads, decreasing the bioavailability of total P loads to Lake Erie and likely constraining cyanoHABs severity (Fig. 1b, Fig. 4a and 4b). This conservative estimate does not include P sorption by suspended sediment retained and buried in the river system or its floodplain. The percent of DRP sorbed was 2.8-fold higher before than after 2003 (< 2003 = 91% ± 0.75%; ≥ 2003 = 33% ± 0.08%; Fig. 4b), a threshold year for increased flow, DRP loading, and cyanoHABs in Lake Erie (*21, 27*).

**Figure 4.**
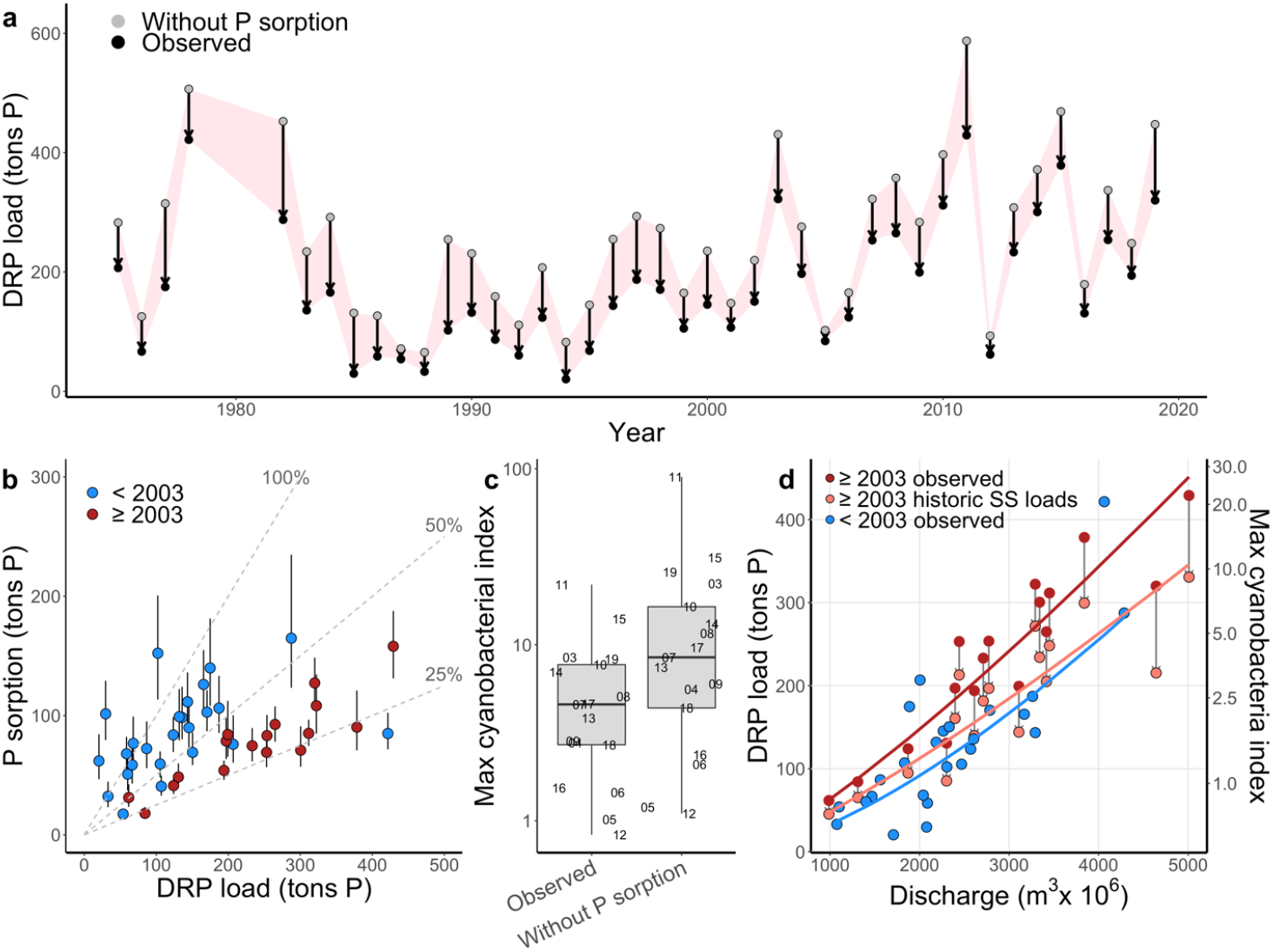
Dynamics, effects, and controls of P sorption in the Maumee River network. **a,** Patterns in Maumee River DRP loads as observed and estimated in the absence of P sorption. **b**, Relationship between P sorption and DRP load illustrating differences in the percent of DRP sorbed (grey lines) for periods before and after 2003. **c**, Estimated effect of P sorption on the maximum cyanobacterial density. **d**, DRP load – discharge relationship before and after 2003 as well as with historical suspended sediment loads. Relationships for before 2003 and after 2003 with historic suspended sediment loads were statistically similar (Table S3), Values in **a-d** were calculated for high flows (<25% flow exceedance) between March and June.

Thus, in support of our hypothesis, P sorption substantially reduces DRP loading to Lake Erie, reducing the bioavailability of total P loads. DRP is entirely bioavailable to cyanoHABs, whereas, only ~26% of Maumee River suspended sediment P is bioavailable (*3*). Furthermore, in this system, immobilization of DRP onto suspended sediment currently represents a long-term P sink because ~70% of sediment exports are buried near the Maumee River mouth (*28*) where bottom sediment have low internal P loading (*29*).

Our results indicate that P sorption in the Maumee watershed performs an unrecognized ecosystem service that constrained cyanoHABs and the resulting ecological, economic, and human health consequences. To estimate the effect of P sorption on cyanoHABs, we predicted (following Stumpf et al. (*21*)) the maximum annual cyanobacteria density in Lake Erie for each year between 2003 and 2019 using March-June DRP loads during high flows estimated with and without P sorption in the river. Without P sorption, the maximum annual cyanobacteria density would be 2.6-fold higher (Fig 4c). Cyanobacteria density estimates in the absence of P sorption during some years were 400% higher than observed in Lake Erie since 2003. While nitrogen or light limitation might keep cyanobacteria from reaching these densities (*30*), riverine P cycles clearly reduce the bioavailability of P exports to Lake Erie and serve as a substantial and previously unknown constraint on cyanoHABs.

Our results indicate that, after accounting for discharge, recent increases in DRP exports to Lake Erie from the Maumee River can be explained by declines in P sorption, driven by declines in suspended sediment loads. After accounting for variation in discharge, which has been increasing for the last 25 years (Fig. S6a, also see (*31*)), Maumee River March–June suspended sediment loads have declined between 1975 and 2019 (Pyear < 0.001; R^2^ = 0.672; Fig. S6b; also see (*31*)). Declines in suspended sediment loads are likely due to changes in agricultural practices that reduce erosion such as the use of cover crops, no-till cropping systems, and tile drains (*27, 32*). Between 1975 and 2019, March–June suspended sediment loads at high discharge (4000 m^3^ x 10^6^) have declined by 48% or 120 kilotons dry mass per decade, resulting in a 48% decline in P sorption (20 tons P per decade). To examine how declines in suspended sediment loads and P sorption contributed to differences in DRP loads before and after 2003, we estimated DRP loads in the absence of historic declines in suspended sediment loads. After accounting for P sorption not occurring due to changes in suspended sediment loads, DRP load – discharge relationships were similar before and after 2003 (P = 0.26; R^2^ = 0.65, Fig. 4d; Table S4).

Thus, recent increases in DRP loads to Lake Erie can be attributed to both increasing discharge and a decline in P sorption associated with declining suspended sediment loads. The impact of other mechanisms for increased DRP loading (e.g., soil stratification, macropores, legacy P, or tile drains) are not well supported empirically at the watershed scale (*27, 33*), but likely also contribute to patterns in discharge and DRP loads. Our results clarify the role stream and river P cycles play in controlling linkages among climate, agricultural practices, and downstream water quality (Fig. 1b).

Climate change may have unanticipated effects on the role river P cycles play in watershed nutrient budgets. In the Maumee watershed, climate change, which is predicted to increase spring precipitation by up to 20% by 2100 (*34*), could increase river discharge (*35*) (but see (*36*)) likely leading to more erosion, higher suspended sediment loads, and more P sorption; potentially decreasing the bioavailability of P loads to Lake Erie. However, the cumulative impact of river P cycling on cyanoHABs will depend upon the impact of rivers on the bioavailability of P exports (Fig. 1b) as well as other climate impacts on agricultural fields and internal P loading in western Lake Erie. While benthic sediment in western Lake Erie currently is a long-term P sink (*29*), changes in water temperature could flip this P sink to a source (*37*).

### Implications for P management

Here, we identify a critical ecosystem service –river P cycling during high flows (Fig. 1b)– that, with changing agricultural practices and climate, helps explain the proliferation of cyanoHABs in Lake Erie and perhaps worldwide. Our findings are broadly applicable to other watersheds with intensive agricultural land use (*6, 38*). Phosphorus sorption-desorption is likely important during high flows in river basins with headwater to lake travel times greater than 12 hours (*13, 14, 39*). In many of these river basins, suspended sediment loads have declined over the last 50 years (e.g., Mississippi, Ohio, Yangtze, and Yellow rivers), due to changes in climate, reservoir impoundment, and agricultural practices (*40–42*). Yet, depending upon whether suspended sediment sorb or release P (an expanding DRP arrow in Fig. 1b), riverine P cycles could enhance or mitigate cyanoHAB severity. Because riverine processes can play a central role in watershed P budgets, as also shown for river carbon and nitrogen cycles (*43, 44*), these processes must be considered as a part of cyanoHAB mitigation and management.

Unfortunately, the role of riverine processes in shaping P exports is rarely considered in watershed scale P studies, watershed models, or P management (*6, 45, 46*). Our work on P cycling during high flows, contributes to numerous low-flow studies demonstrating how river P cycles can alter the magnitude and bioavailability of P exports (*7, 47*). Failing to consider the contribution of river P cycles to downstream exports can result in a misattribution of P sources or sinks to the wrong location or mechanism (Fig. 1). Thus, not including rivers in watershed models and P management ignores an important watershed scale process influencing the bioavailability of P loads and cyanoHABs in recipient ecosystems.

Our research also contributes to our growing understanding of the consequences of managing for a single stressor or constituent (*48, 49*). We demonstrate that while efforts to reduce sediment erosion in the Maumee River watershed were successful, declines in suspended sediment loads contributed to increases in the bioavailability of P loads and likely cyanoHABs. Our results do not indicate that efforts to reduce erosion and sedimentation should be halted. Such practices provide benefits to agricultural systems (*19*) and reduce sediment pollution in freshwater ecosystems (*50*). Instead, our research emphasizes the need to simultaneously consider multiple stressors with a watershed-scale approach (*38*). This approach considers and addresses potential unanticipated consequences which could undermine management, waste conservation funds, and erode public trust.

## Acknowledgments

We thank S. Trail and J. Vann for help in the field and laboratory as well as C. Dolph, J. Finlay, L. Johnson, E. Marschall, and T. Williamson for comments on an early draft which improved this work. We also thank L. Johnson for help identifying sampling sites and water quality monitoring data.

## Author contributions

Conceptualization: WMK and JMH

Methodology: WMK, JMH, and SC

Investigation: WMK, JMH, and SC

Formal Analysis: WMK and JMH

Visualization: WMK and JMH

Funding acquisition: WMK and JMH

Project administration: JMH

Supervision: JMH

Writing – original draft: WMK and JMH

Writing – review & editing: WMK, JMH, and SC

## Data and materials availability

All data and analysis scripts can be found in our GitHub repository: https://github.com/hood211/HighFlowSorption.git.

## Supplementary Materials for

### Materials and Methods

#### Stormwater sampling

Our six focal stream sites were located within the Maumee River watershed (17,000 km^2^; Fig. 1c; Table S1) which is dominated by agricultural land use (~87%) and contains glacial till or lake plane soils with poor natural drainage (*51*). We sampled our focal streams between January and June 2019 for suspended sediment concentration, dissolved reactive P (DRP), and aspects of P sorption-desorption (sorption isotherms as well as sorption rates). Each site was located near a USGS stream gauging station. Four sampling sites cooccurred with a Heidelberg University National Center for Water Quality Research (NCWQR) water quality monitoring station (Fig. 1c).

Each focal stream was sampled once per storm event (Fig. S1) from a bridge by deploying 19 L containers to collect 75 L of stream water from the center of the main channel. Stream water was stored in these containers until processed in the laboratory.

Immediately following the collection of stream water, we homogenized the sample and collected triplicate dissolved reactive phosphorus and suspended sediment samples (expressed in terms of dry mass [DM]). DRP samples were immediately filtered through 0.45 μm glass fiber filters (Environmental Express, Charleston, SC, USA) into 20 mL HDPE scintillation vials (Fisher Scientific, Pittsburgh, PA, USA) using acid-washed syringes and kept cold (~ 4°C) until frozen in the laboratory. Dissolved reactive phosphorus was analyzed within two days of collection on a spectrophotometer using the colorimetric molybdenum blue reaction method (*52*). Suspended sediment samples were stored in 250 mL acid-washed HDPE plastic bottles, which were kept cold (~ 4°C) until filtered onto pre-weighed and ashed glass fiber filters (Whatman^TM^ GF/F) within 24-hours. Sediment DM filters were dried at 60 °C for at least 48 hours and then reweighed.

The remainder of stream water was used to measure aspects of sediment P sorption-desorption. This water was transported to the laboratory in 19 L containers and stored in a dark environmental chamber set to the mean water temperature of the six streams. To consolidate the suspended sediment for analyses, we used a WVO Raw Power Continuous Flow Centrifuge (6000 RPM; WVO designs, Charleston, SC, USA). Stream water was transferred to a 20 L Nalgene carboy and set to a low flow-through rate (~2 L h^−1^) to minimize sediment loss through the centrifuge outflow. Sorption rate and isotherm measurements were conducted within one week.

#### P sorption-desorption

To characterize suspended sediment – DRP interactions, we used a multi-tiered approach which began with measurement of P sorption-desorption isotherms and ended with the calculation of mass-specific (μg P mg DM^−1^ h^−1^) and volumetric (μg P L^−1^ h^−1^) P sorption-desorption. We calculated several aspects of P sorption-desorption from P sorption isotherms including: the zero-equilibrium phosphorus concentration (EPC_0_; μg P L^−1^), the P sorbed to suspended sediment particles at the time of collection (S_amb_; ug P mg DM^−1^), the maximum P suspended sediment particles could sorb (S_max_; ug P mg DM^−1^), and the capacity for P sorption by suspended sediment particles during downstream transport (Scap; ug P mg DM^−1^).

##### P sorption isotherms

We measured P sorption-desorption isotherms following Jarvie et al. (*53*). Briefly, in 50 mL Falcon centrifuge tubes (Thermo Fisher Scientific, Corning, NY, USA), we combined consolidated suspended sediments (~ 0.1-0.2 g) from each stream with an ‘artificial river water matrix’ similar in conductivity and calcium concentration to river water. To create this matrix, we mixed CaCl2 and deionized water at concentrations sufficient to mimic the specific conductivity of the Maumee river at noon on the day of sampling (Waterville, OH; USGS station number: 04193490). Next, we spiked the centrifuge tubes with a 1000 μmol P stock solution (KH_2_PO_4_) to create a DRP concentration gradient (0, 2.5, 5, 10, 15, 20 μmol P). Then, we placed the centrifuge tubes on an orbital shaker at 150 RPM in a dark environmental chamber set to the mean water temperature of the six streams. After 24 hours, we stopped the reaction by centrifuging the solution at 5000 RPM for 10 min and immediately filtered the supernatant through 0.45 μm glass fiber filters (Environmental Express, Charleston, SC, USA) using acid-washed syringes into 20 mL HDPE scintillation vials (Fisher Scientific, Pittsburgh, PA, USA). Samples were frozen until analyzed for DRP. The sediment in each centrifuge tube was dried at 70 °C for 48 hours and weighed.

To estimate P sorption parameters, we fit data obtained from the P sorption isotherm experiments to a Langmuir adsorption isotherm model following Lai and Lam (*54*). We selected the Langmuir model because it was a good fit to our data and provided parameters relevant to subsequent analyses (e.g., *S*_amb_ and *S*_max_). In the Langmuir isotherm model, DRP removed during the 24 h incubation (*S*_24_) is

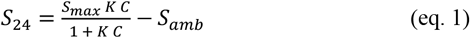

where, *K* is the bonding energy constant, and *C* is the DRP concentration of the supernatant after the 24-hour incubation. Sorption capacity (*S*_cap_) is the difference between *S*_max_ and *S*_amb_. Our model fits were evaluated based on visual evaluation of residuals and model residual standard errors. One or two outliers were removed from twelve out of 78 isotherm models. Data from four sorption isotherms did not conform to the Langmuir model. Of those, three were fit with a linear isotherm model relating *S*_24_ and *C* while one isotherm did not conform to any model we evaluated.

We used the Langmuir model parameters to estimate *EPC*_0_, the DRP concentration which, when placed in contact with sediment, produces no net exchange of DRP in solution over a 24-hour period. We calculated *EPC*_0_ by rearranging equation 1:

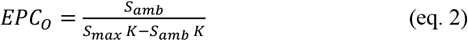

*EPC*_0_ can be used to characterize whether sediments sorb or desorb P. When *EPC*_0_ is greater than stream DRP concentrations sediments release DRP, whereas, sediments sorb DRP when *EPC*_0_ is less than stream water DRP concentrations (*55*).

##### P sorption-desorption rates

We measured P sorption and desorption rates by suspended sediments following Jarvie et al (*53*); however, since suspended sediments desorbed P under ambient conditions in only one of 78 samples (based on the difference between *EPC*_0_ and stream water DRP concentrations) we only present P sorption rates. To measure P sorption rates, we added 0.1 – 0.2 g of consolidated suspended sediments to 50 mL Falcon centrifuge tubes containing an ‘artificial river water matrix’ (see above) enriched with a P stock (as KH_2_PO_4_) to the stream water DRP concentration at the time of sampling. We shook the cepended sediments to 50 mL Falcon centrifuge tubes containing an ‘artificial river water matrix’ (see abontrifuge tubes on an orbital shaker (set at 150 RPM) located in a dark environmental chamber, which was set to the mean stream temperature at the time of sampling, for the following incubation periods: 0, 1, 3, 5, 10, 15, 30, 45, and 60 minutes. Pilot experiments indicated that DRP concentrations reached equilibrium within less than one hour. To terminate the incubation, we centrifuged the samples and collected DRP and sediment samples as described above.

We calculated suspended sediment P sorption on a dry mass-specific (*S*_S,MS_, μg P g DM^−1^ h^−1^) and volumetric basis (*S*_S,V_, μg P L^−1^ h^−1^). First, we estimated the proportional loss rate (*k*) as the slope of the relationship between log-transformed DRP per sediment mass (μg P g DM^−1^) and time. Prior to fitting this model with simple linear regression, we removed samples representing time points after DRP concentrations converged to an equilibrium since these data did not meet model assumptions of log-linear change over time. Mass-specific P sorption rate is

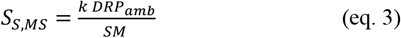

where *DRP*_amb_ is the ambient stream water DRP concentration (μg DRP L^−1^) and *SM* is the mean sediment dry mass (g DM) in the centrifuge tubes. Volumetric P sorption (*S*_S,V_) in the stream at the time of sediment collection is

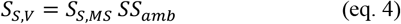

where *SS*_amb_ is the stream water total suspended solid concentration (g DM L ^−1^) at the time of sediment collection in each stream.

#### Discharge and flow exceedance

Discharge data for each tributary and the Maumee River was retrieved from National Water Information System (http://waterdata.usgs.gov/nwis) (*56*). We used the available data for each system (Table S1) to calculate the percent of time each flow was exceeded during the record as:

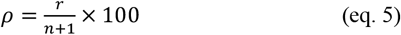

where *r* is the rank of the 15-minute flow according to its magnitude and *n* is the total number of 15-minute flows in the tributaries or water quality samples for the Maumee River (*57*).

#### Total P sorption in tributaries

To estimate the impact of P sorption on DRP export from our focal tributaries at the watershed scale, we combined our P sorption estimates with publicly available discharge, DRP, and suspended sediment data to estimate P sorption by all sediments transported by our sampling sites between March and June 2019 at 0-25% flow exceedance (hereafter high flow). Only three of our six focal tributaries (STF, UTLC, and WC) had sufficient DRP and suspended sediment data to support this analysis. To begin, we combined NCWQR (*58*) water quality data with USGS discharge data to estimate loading rates. Since NCWQR water quality data was collected less frequently (~1-3x per day) than the USGS discharge data (~every 15 minutes), we used generalized additive mixed models (GAMM) to predict the missing DRP and suspended sediment concentrations (*59*). Our GAMMs for DRP and suspended sediment concentrations used a thin plate regression spline for discharge and a cyclic cubic regression spline for month. All models had normal residuals, little prediction bias, and high R^2^’s (R^2^_adj_ = 0.46 – 0.58). GAMMs were fit with the “gamm” function in the package “mgcv” (*60*) in the R statistical environment (*61*). We calculated DRP and suspended sediment loads as the product of discharge, the time between discharge measurements (generally 15 minutes), and DRP or suspended sediment concentrations, respectively.

Next, we combined water quality and discharge data with sorption rate measurements to estimate P sorption by suspended sediments during high flow events between March and June 2019. Our approach accounted for both the P sorption capacity of suspended sediments and the influence of DRP concentrations on P sorption-desorption by sediments. We began by estimating a watershed-scale mass-specific sorption rate (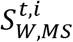; g P g DM^−1^ h^−1^) in tributary *t* for each time *i* (associated with a discharge measurement) as:

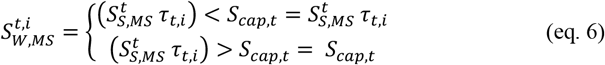

where *τ_t,i_* is travel time in tributary *t* at time *i* (h; described below), 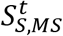 is the mass-specific P sorption of sediment in tributary *t*, and potential watershed-scale mass-specific sorption 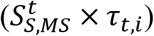 is constrained to be no greater than the P sorption capacity of the suspended sediments in tributary *t* (*S*_cap,t_;g P g DM^−1^). Then, we calculated watershed-scale volumetric sorption (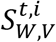; g P g m^−3^ h^−1^) in tributary *t* at time *i* as:

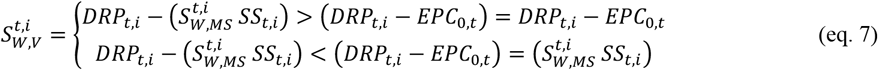

where *DRP*_t,i_ (g P m^−3^) and *SS*_t,i_ (g DM m^−3^) are the DRP and suspended sediment concentrations in tributary *t* at time *i*, respectively, and potential watershed-scale volumetric sorption 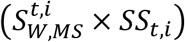 is constrained so that P sorption does not reduce DRP concentrations below *EPC*_0_, which reflects an equilibrium between sediment sorption-desorption and stream water DRP. Finally, we estimated total P sorption (*S*_Tot,t_; metric ton P) in tributary *t* integrating across all time periods meeting our criteria (i.e., all *i*’s) as:

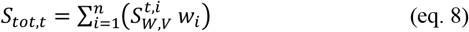

where *w_i_* is the time between discharge measurements (i.e., times *i-1* to *i*). To estimate the uncertainty associated with our measurements, we bootstrapped these calculations (n = 500) using all estimates of sorption rate, sorption capacity, and *EPC*_0_ for tributary *t*.

#### Total P sorption in Maumee River network

To estimate the total net P sorption within the Maumee River network at 25 percent exceedance flows between March and June in each year from 1975 to 2019, we combined watershed-scale mass-specific P sorption from the three tributaries 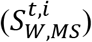, with publicly available discharge, DRP, and suspended sediment data for the Maumee River at Waterville, OH (retrieved from NCWQR (*58*)). Maumee River sediment and DRP loads were calculated as the product of DRP or suspended sediment concentration, discharge, and the time between samples. We calculated Maumee River mass-specific P sorption for time period *w_i_* as:

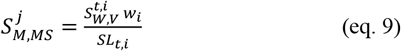

where *SL_t,i_* is the sediment load in tributary *t* during the time period *w_i_*. Maumee River mass-specific P sorption incorporates watershed-scale constraints on P sorption including: P sorption capacity, the influence of DRP on P sorption-desorption dynamics, and the capacity of sediments to sorb P during downstream transport to the monitoring site on the Maumee River at Waterville, OH. Total P sorption by Maumee River suspended sediment during periods meeting our criteria in year *y* is:

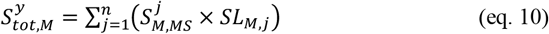

where SL_M,j_ is the Maumee River suspended sediment load during time window *w_i_*. To estimate the uncertainty associated with total P sorption we bootstrapped these calculations (n = 500) using all estimates of watershed-scale mass-specific sorption rate 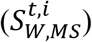 from the tributaries.

#### Impact of sorption on cyanoHABs

Finally, to determine the impact of P sorption on cyanoHABs in western Lake Erie, we estimated the maximum cyanobacterial index (*CI*_max_; 10^20^ cells), a satellite-based measure of the annual maximum cyanoHABs density, for each year between 2003 and 2019 following Stumpf et al. (*21*):

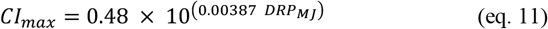

where DRP_MJ_ is the DRP load (metric tons) for periods between March and June with flows greater than 25% exceedance. Since this model was calibrated using data from 2002 to 2015, we did not estimate maximum cyanobacterial index for years before 2002. This model provides a good fit to the data; however, a more widely used model which uses total bioreactive P and includes July loads provides a slightly better fit for some years (*21*). We used equation 10, instead of other alternatives, because it was simple and did not require extrapolating P sorption beyond our sampling period.

#### Travel time

For each reach between stream gauging stations, we estimated travel time (*τ*) following Du et al. (*62*):

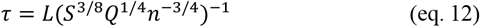

where *L* is the reach length between gauging stations (km), *S* is the mean river slope between gauging station (*km km*^−1^), *Q* is discharge (*m*^3^*min*^−1^), and n is Manning’s coefficient (*m*^1/3^ *min*^−1^). Reach lengths and slopes between our six tributary sampling sites and the Maumee River at Waterville, OH (Fig. S2) were calculated using the qProf plugin in QGIS (Version 3.8). For the Manning’s coefficient, we selected a value of 6.2*x*10^−3^ *m*^1/3^ *min*^−1^ which was estimated for the Middle Fork of the Vermilion River (near Danville, IL) (*63*). This site was similar to our tributaries in terms of riverbed and bank composition.

#### Statistical analyses

To examine how discharge and stream identity influences tributary P sorption and suspended sediment loads, we used linear models and an information theoretic approach (*64*). Models were ranked by AICC and the models with a delta AICC less than two were considered to have the most support (*64*). To examine the relationship between tributary sediment equilibrium phosphorus concentration (EPC_0_) and DRP concentration, we used standardized major axis regression using the *smatr* package (*65*). We selected this approach over a general linear model because we wished to estimate the slope of the EPC_0_-DRP relationship and because, due to a poor understanding of whether EPC_0_ was shaping DRP or vice versa, we considered these regression terms symmetric. We also used linear models to examine how Maumee River DRP load – discharge relationships changed over time. In this case, we compared p-values, following the guidelines of Wasserstein (*66*), for a grouping variable separating 1975–2002 and 2003–2019.

To examine the influence of declines in suspended sediment loads between 1975 and 2019 on P sorption and ultimately DRP loading to Lake Erie, we had to account for intra-annual changes in discharge. We accomplished this by examining changes in the DRP load–discharge relationship, following a four-step process. We built a linear model relating Maumee River high flow March–June suspended sediment loads to discharge and year. We used that model to predict suspended sediment loads for each year between 1975 and 2019 with year in the prediction equation fixed to 1975. We used these predicted suspended sediment loads, which account for interannual changes in discharge, to estimate P sorption (approach described above) and, finally, DRP loads with historic (i.e., 1975) suspended sediment loads and P sorption. We used linear models to examine how the DRP load–discharge relationships differed between 1975 to 2002 observed DRP loads and 2003 to 2019 DRP loads predicted with historic suspended sediment loads and P sorption rates. Prior to analyses, we used qqplots to assess normality assumptions. Most terms were log_10_ transformed while flow exceedance was raised to the 6^th^ power for analyses. All data and analysis scripts can be found in our GitHub repository: https://github.com/hood211/HighFlowSorption.git.

## Supplementary Figures

**Figure S1.**
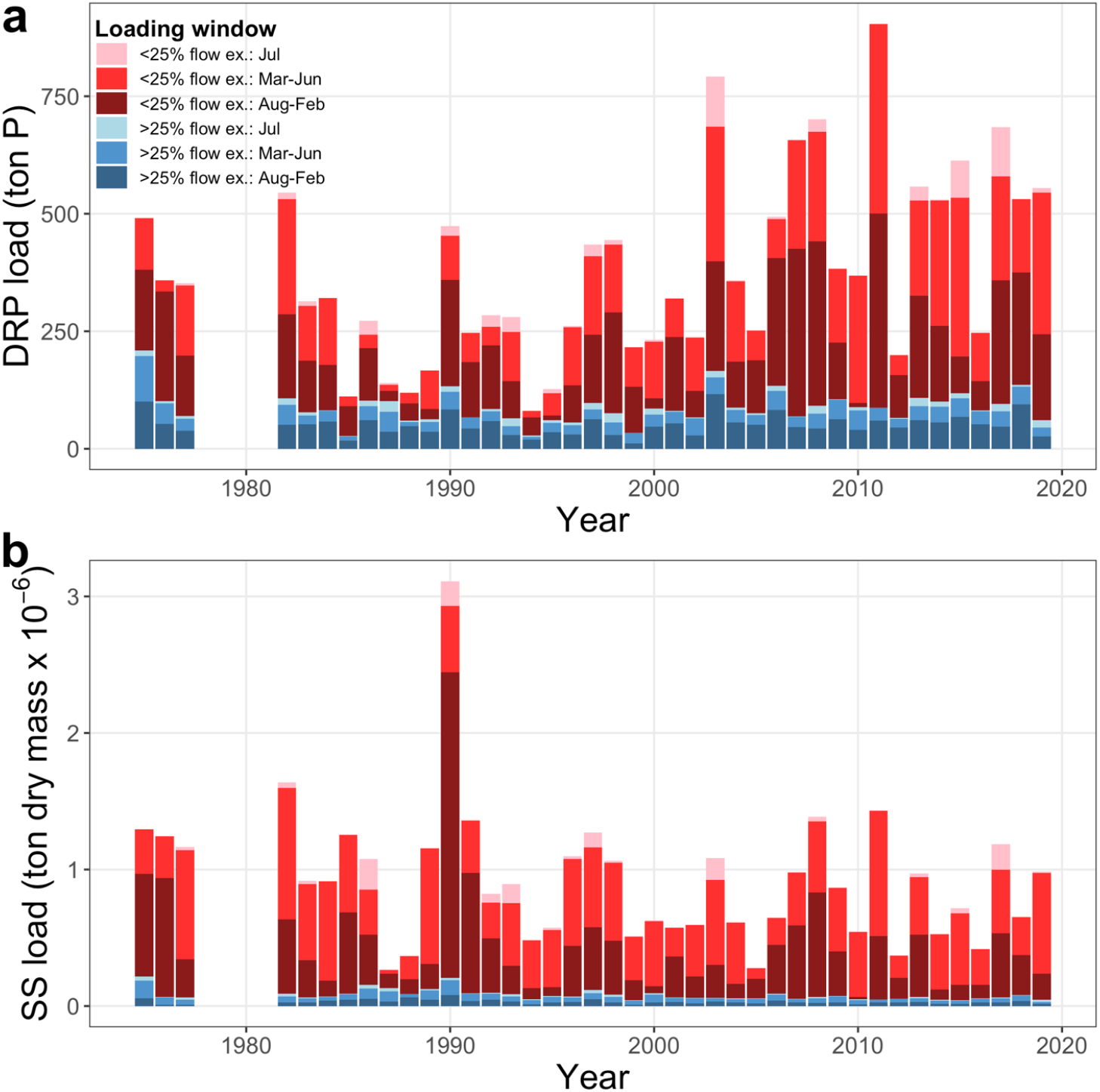
Dissolved reactive phosphorus (DRP, **a**) and suspended sediment (SS, **b**) loads during low (>25% flow exceedance) and high (<25% flow exceedance) flows integrated across three time windows: August–February, March–June, and July. In contrast to Lake Erie cyanoHAB forecast models, this study focuses on high flow periods during March–June because that window overlapped with our tributary sampling and because July represented a small portion of March-July DRP (7.9% ± 12.3%; mean of annual estimates between 1975 and 2019 ± 1 standard deviation) and suspended sediment loads (5.6% ± 10.0%). Between March and June, high flow DRP and suspended sediment loads represent 75.0% (± 14.8%) and 88.0% (±12%), respectively of total loads during these time windows. Note that sufficient data was not available during 1978-1981 to calculate loads.

**Figure S2.**
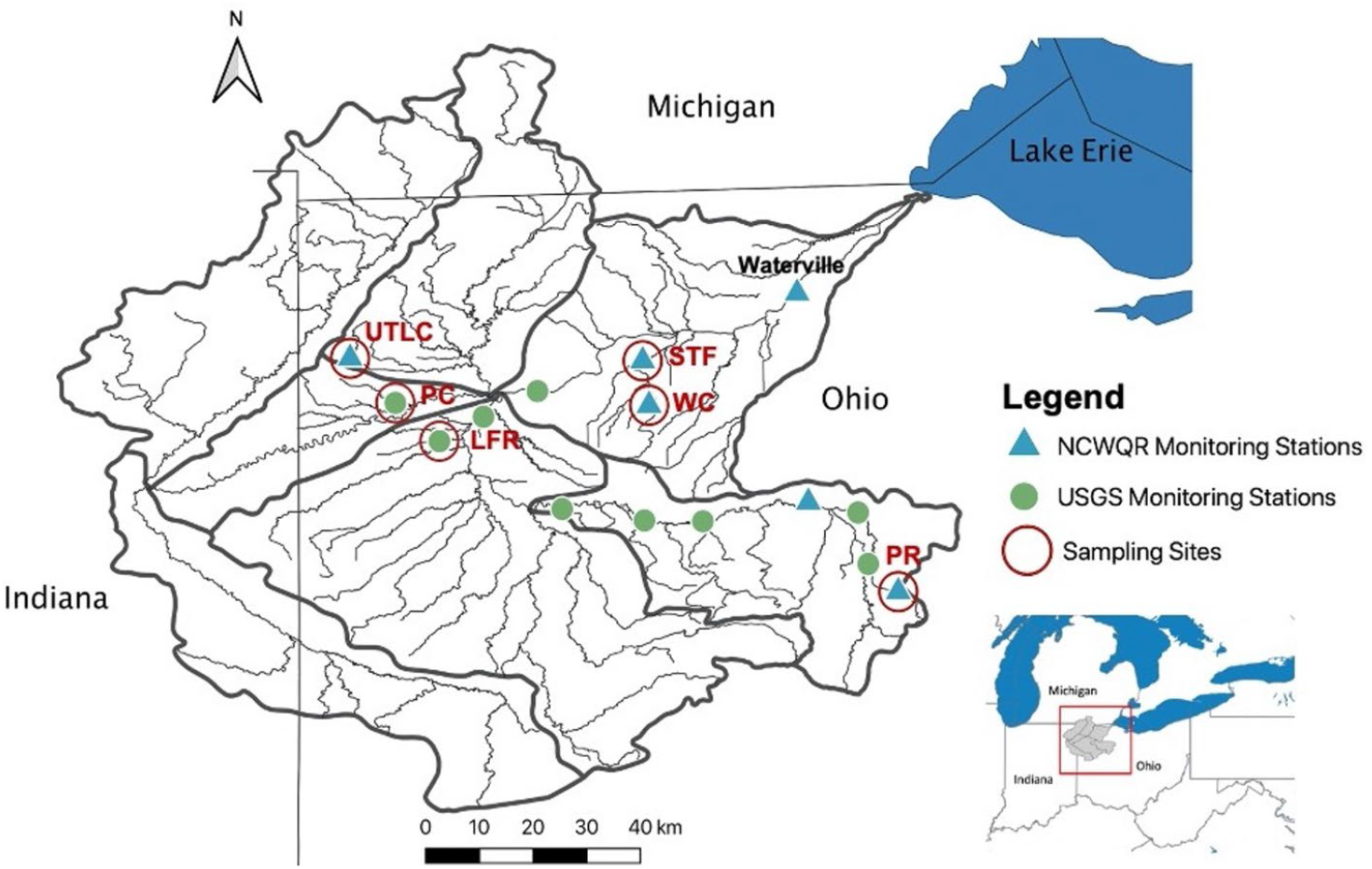
Map of the Maumee River watershed illustrating the location of study sites and relevant discharge (U.S. Geological Survey [USGS]) and water quality (National Center for Water Quality Research [NCWQR]) monitoring stations.

**Figure S3.**
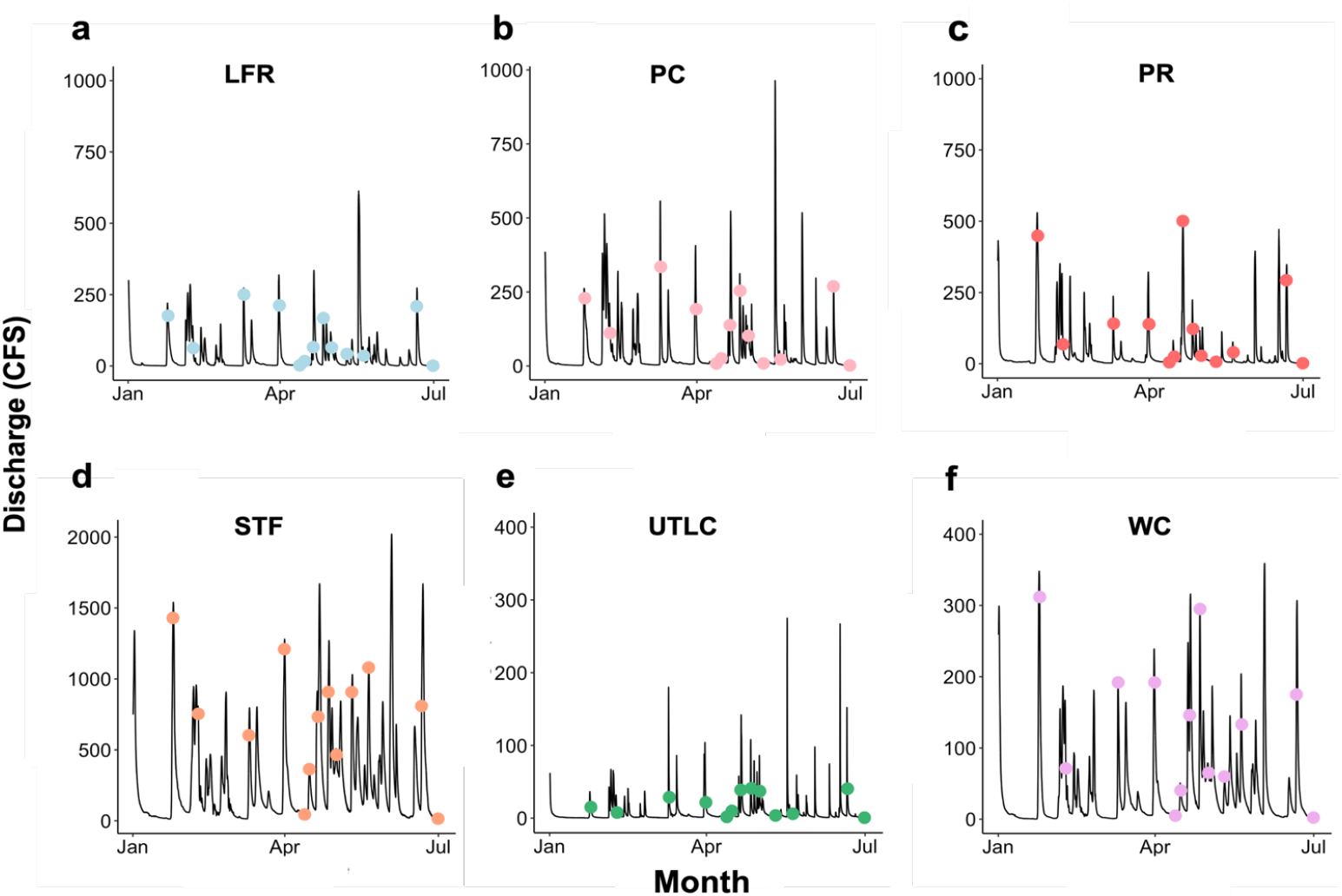
Hydrograph of the six focal streams from January to June 2019 (**a-f**). Points indicating timing of instream sampling. The y-axis scale differs among hydrographs.

**Figure S4.**
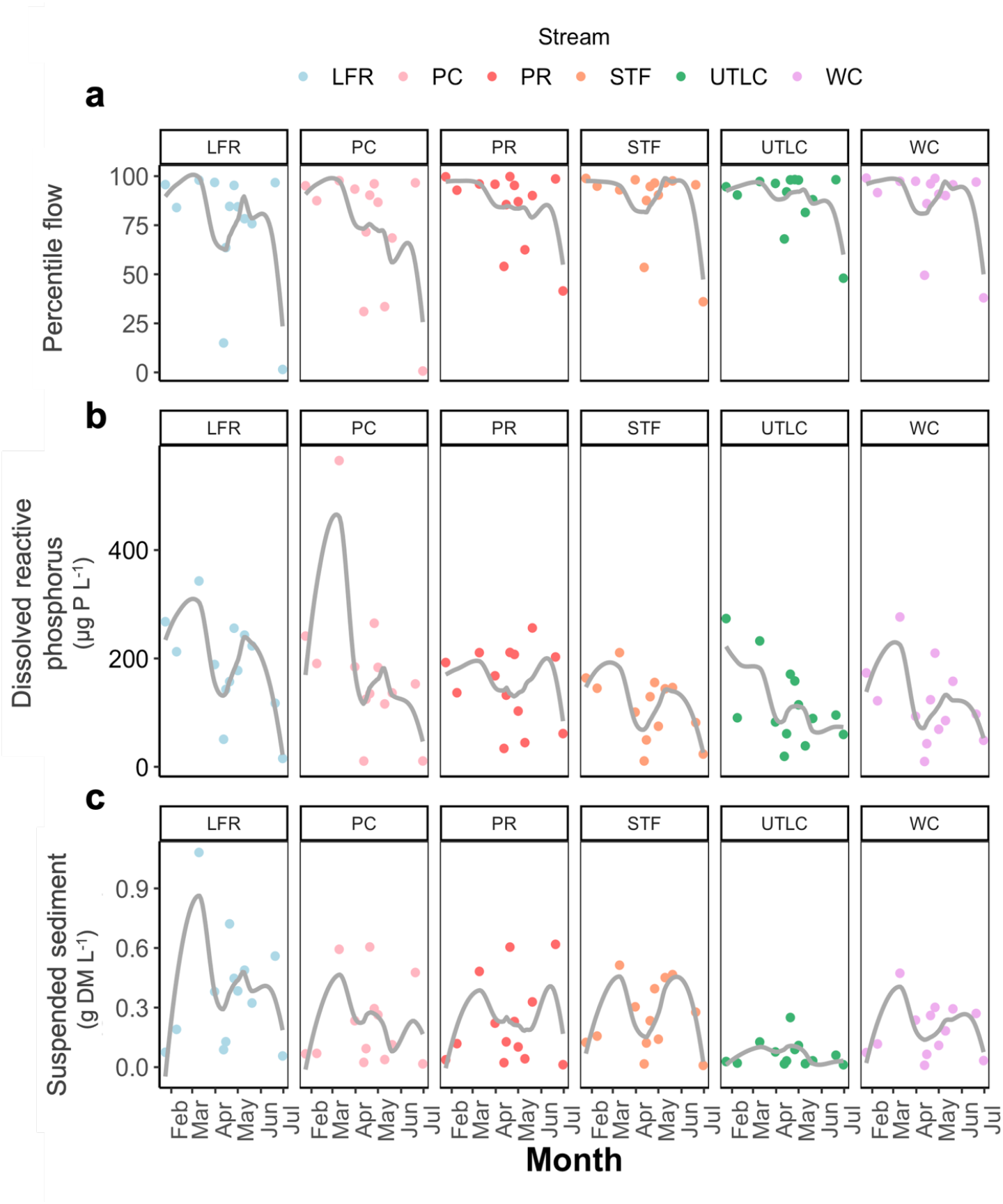
Percentile flow (**a**), dissolve reactive phosphorus (DRP; **b**), and suspended sediment dry mass (DM; **c**) for all stream sites between January and June 2019. Grey lines are loess fits.

**Figure S5.**
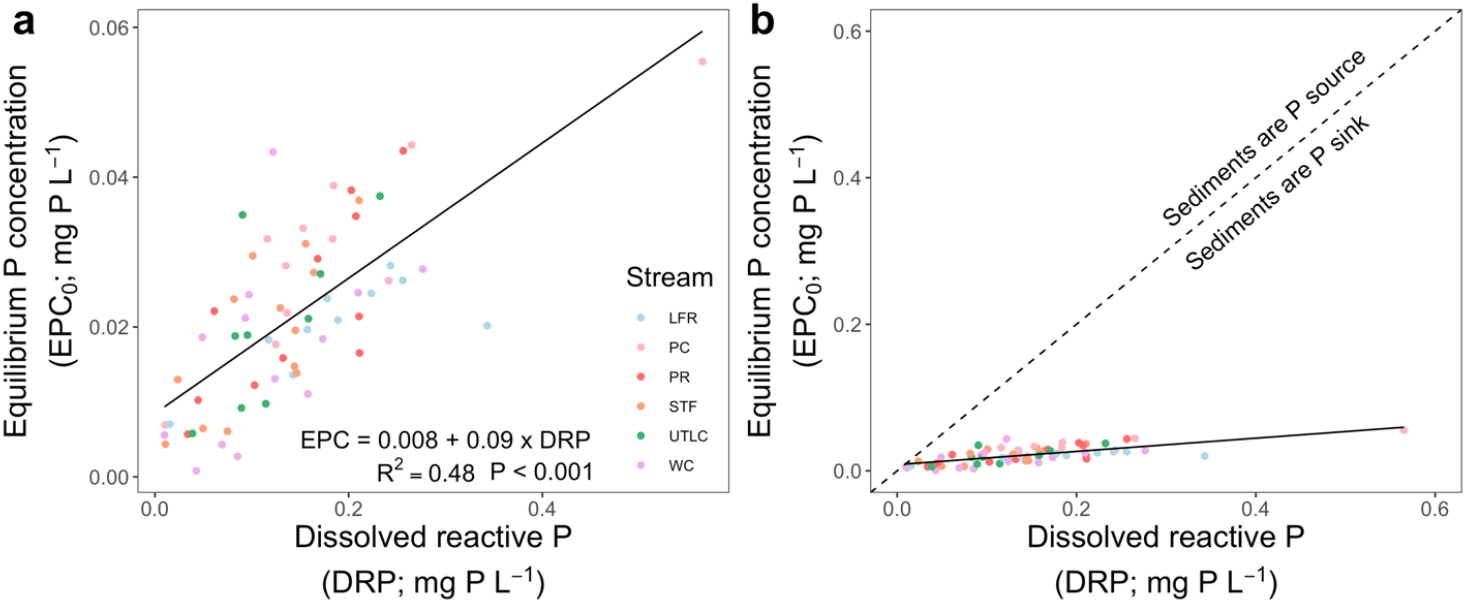
Relationships between suspended sediment equilibrium phosphorus concentration (EPC_0_) and stream water dissolved reactive phosphorus (DRP) concentrations in six Maumee River tributaries with major standardized axis model regression line and statistics (**a**). These same data are also presented with equal x- and y-axis scales as well as a one-to-one line (dashed line) to illustrate P source/sink behavior (**b**). Sediment samples with values below the one-to-one line are P sinks while those with values above the one-to-one line are P sources.

**Figure S6.**
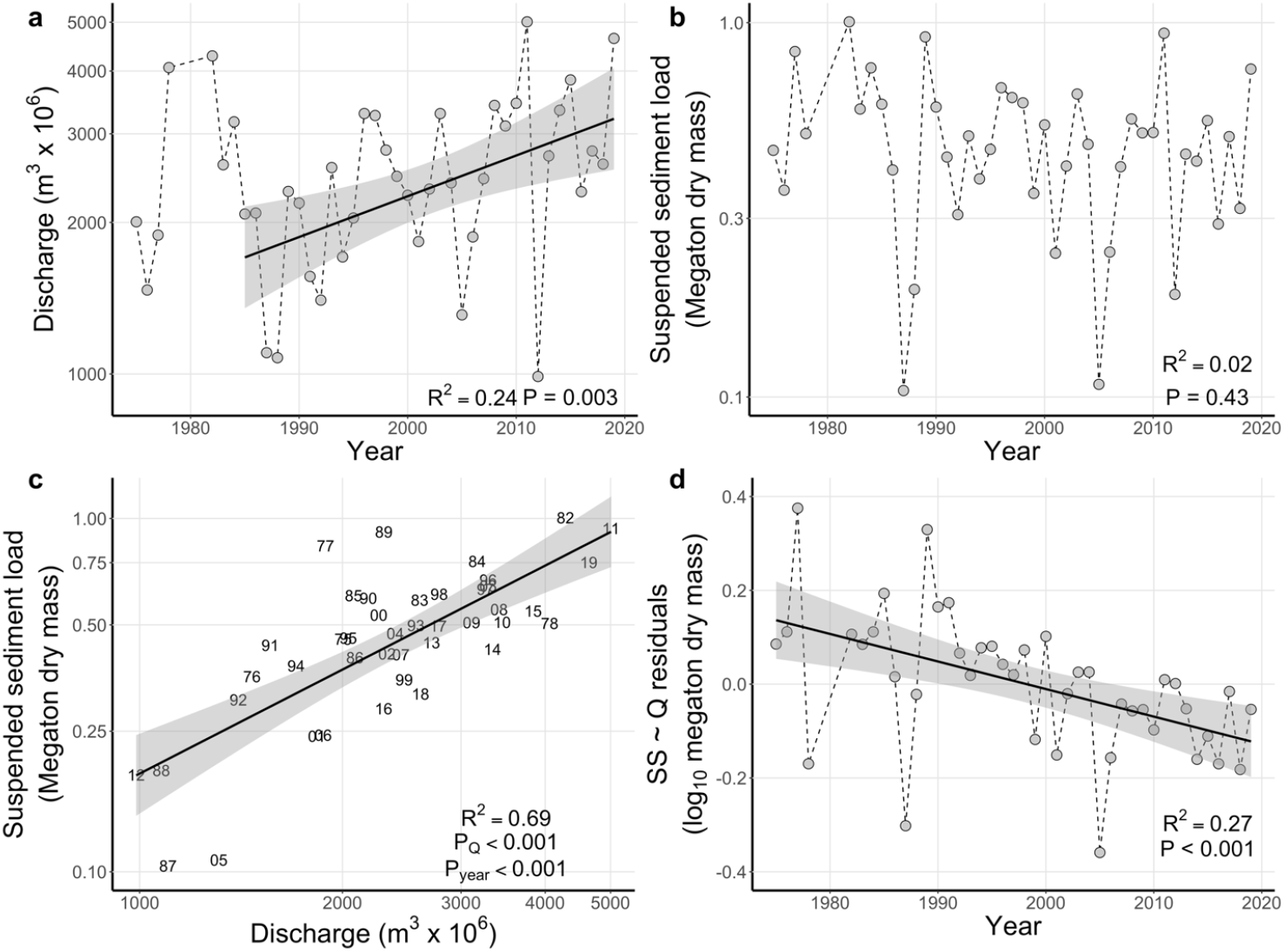
In the Maumee River, discharge has generally increased over time since 1985 (**a**), whereas suspended sediment loads have not changed (**b**). However, suspended sediment loads increase with discharge (**c**) and, when discharge is held constant (in a multiple regression model) suspended sediment loads have decreased between 1975 and 2019 (**d**). Panel **d** shows the residuals of the suspended sediment (SS) – discharge (Q) relationship and year. All values are for high flows (<25% flow exceedance) between March and June. Regression statistics are from general linear models. Panel **c** provides R^2^ and p-values from a model relating suspended sediment loads to discharge and time whereas panel **d** provides regression statistics for a linear regression between SS – Q residuals and year.

## Supplementary Tables

**Table S1.** Site information for the six focal tributaries and the Maumee River including stream code, USGS site IDs, drainage area stream order based on the Strahler method, and the years of discharge data used to calculate maximum flow.

| Stream name | Stream code | USGS Site ID | Drainage area (km<sup>2</sup>) | Stream order (Strahler) | Years used to calculate flow exceedance |
| --- | --- | --- | --- | --- | --- |
| Little Flat Rock | LFR | 04191444 | 39.6 | 2 | 2015-2019 |
| Platter Creek | PC | 04183979 | 50.5 | 2 | 2018-2019 |
| Potato Run | PR | 04188324 | 44.5 | 1 | 2017-2019 |
| South Turkey Foot | STF | 04192599 | 300.4 | 3 | 2015-2019 |
| Unnamed Tributary to Lost Creek | UTLC | 04185440 | 10.8 | 1 | 2015-2019 |
| West Creek | WC | 04192574 | 40.2 | 1 | 2015-2019 |
| Maumee River at Waterville, OH | MR | 04193500 | 17,000 | 7 | 1987-2019 |

**Table S2.** Model selection table examining how tributary suspended sediment load (SS) and P sorption are shaped by discharge (expressed as flow exceedance probability), tributary identity, and suspended sediment loads.

| **Model** | **df** | **AIC<sub>c</sub>** | **ΔAIC<sub>c</sub>** | **Weight** | **R<sup>2</sup>** |
| --- | --- | --- | --- | --- | --- |
| *Suspended sediment load* |  |  |  |  |  |
| ~ flow ex. + tributary | 8 | 136.2 | 0.00 | 0.95 | 0.75 |
| ~ flow ex. × tributary | 13 | 142.2 | 5.94 | 0.05 | 0.78 |
| ~ flow ex. | 3 | 181.8 | 45.61 | 0 | 0.41 |
| *P sorption* |  |  |  |  |  |
| ~ SS + tributary | 8 | 187.8 | 0 | 0.65 | 0.81 |
| ~ SS × tributary | 13 | 189.2 | 1.43 | 0.32 | 0.84 |
| ~ SS | 3 | 194.1 | 6.36 | 0.27 | 0.75 |
| ~ flow ex. + tributary | 8 | 231.9 | 44.12 | 0 | 0.69 |
| ~ flow ex. × tributary | 13 | 234.6 | 46.85 | 0 | 0.63 |
| ~ flow ex. | 3 | 246.5 | 58.68 | 0 | 0.45 |

**Table S3.** Linear regression model tables for relationship between log_10_ DRP load and log_10_ discharge for observed values during two time windows and for DRP loads predicted with historic sediment loads. Values provided for intercept and discharge are the coefficient, standard error, and p-value. P-values less than 0.05 are in bold to aid interpretation. Discharge and DRP load values are for the greater than 25% exceedance flows between March and June. These data are shown in Fig. 4d.

|  | **Observed** |  | **Predicted with historic sediment loads** |
| --- | --- | --- | --- |
| **Time window** | **(1975 – 2002)** | **2003 – 2019** | **2003 – 2019** |
| Intercept | -2.91 (± 0.94; **0.005**) | -1.86 (± 0.34; < **0.001**) | -2.00 (± 0.48; < **0.001**) |
| Discharge | 1.48 (± 0.28; < **0.001**) | 1.22 (± 0.10; < **0.001**) | 1.22 (± 0.14; < **0.001**) |
| Adjusted R<sup>2</sup> | 0.52 | 0.90 | 0.83 |
| df | 23 | 15 | 15 |
| F-statistic | 27.44 | 148 | 76.9 |
| P-value | < 0.001 | < 0.001 | < 0.001 |

**Table S4.** Linear regression model tables for relationship between log_10_ DRP load and log_10_ discharge (Q) supporting two comparisons. The first group (G) examines differences in DRP – discharge relationships between 1975 – 2002 and 2003 – 2019. The second group examines differences in DRP – discharge relationships comparing 1975 – 2002 observed DRP loads with 2003 – 2019 DRP loads predicted with historic sediment loads. Note that for the second comparison the group effect is not significant. Values provided for intercept and discharge are the coefficient, standard error, and p-value. P-values less than 0.05 are in bold to aid interpretation. Discharge and DRP load values are for greater than 25% exceedance flows between March and June. These data are shown in Fig. 4d.

| Group | 1975 – 2002 observed v. 2003 – 2019 observed |  | 1975 – 2002 observed v. 2003 – 2019 predicted with historic sediment loads |  |
| --- | --- | --- | --- | --- |
| Intercept | -2.91 (± 0.76; < **0.001**) | -2.50 (± 0.55; < **0.001**) | -2.91 (± 0.78; < **0.001**) | -2.51 (± 0.56; < **0.001**) |
| Q | 1.48 (± 0.23; < **0.001**) | 1.35 (± 0.16; < **0.001**) | 1.48 (± 0.23; < **0.001**) | 1.36 (± 0.17; < **0.001**) |
| Q + G | 1.06 (± 1.12; 0.35) | 1.18 (± 0.06; **0.002**) | 0.91 (± 1.15; 0.43) | 0.07 (± 0.06; 0.26) |
| Q × G | -0.26 (± 0.33; 0.44) | – | -0.25 (± 0.34; 0.47) | – |
| Adjusted R<sup>2</sup> | 0.71 | 0.71 | 0.64 | 0.65 |
| df | 38 | 39 | 38 | 39 |
| F-statistic | 33.95 | 51.14 | 25.54 | 38.49 |
| P-value | < 0.001 | < 0.001 | < 0.001 | < 0.001 |

## References and Notes

1. H. W. Paerl et al., Mitigating cyanobacterial harmful algal blooms in aquatic ecosystems impacted by climate change and anthropogenic nutrients. Harmful Algae 54, 213–222 (2016).

2. C. R. C. Kouakou, T. G. Poder, Economic impact of harmful algal blooms on human health: a systematic review. J Water Health 17, 499–516 (2019).

3. D. B. Baker et al., Phosphorus loading to Lake Erie from the Maumee, Sandusky and Cuyahoga rivers: The importance of bioavailability. Journal of Great Lakes Research 40, 502–517 (2014).

4. P. Glibert, M. Burford, Globally Changing Nutrient Loads and Harmful Algal Blooms: Recent Advances, New Paradigms, and Continuing Challenges. Oceanography 30, 58–69 (2017).

5. A. N. Sharpley, P. J. Kleinman, P. Jordan, L. Bergstrom, A. L. Allen, Evaluating the success of phosphorus management from field to watershed. J Environ Qual 38, 1981–1988 (2009).

6. H. P. Jarvie et al., Phosphorus mitigation to control river eutrophication: murky waters, inconvenient truths, and “postnormal” science. Journal of Environmental Quality 42, 295–304 (2013).

7. P. J. Withers, H. P. Jarvie, Delivery and cycling of phosphorus in rivers: a review. Science of the Total Environment 400, 379–395 (2008).

8. D. L. Correll, T. E. Jordan, D. E. Weller, Transport of nitrogen and phosphorus from Rhode River watersheds during storm events. Water Resources Research 35, 2513–2521 (1999).

9. L. E. Gentry, M. B. David, T. V. Royer, C. A. Mitchell, K. M. Starks, Phosphorus transport pathways to streams in tile-drained agricultural watersheds. J Environ Qual 36, 408–415 (2007).

10. A. N. Sharpley et al., Phosphorus loss from an agricultural watershed as a function of storm size. Journal of Environmental Quality 37, 362–368 (2008).

11. A. C. Edwards, P. J. A. Withers, Transport and delivery of suspended solids, nitrogen and phosphorus from various sources to freshwaters in the UK. Journal of Hydrology 350, 144–153 (2008).

12. P. A. Bukaveckas, Effects of channel restoration on water velocity, transient storage, and nutrient uptake in a channelized stream. Environmental Science and Technology 41, 1570–1576 (2007).

13. F. H. Verhoff, D. A. Melfi, S. M. Yaksich, Storm travel distance calculations for total phosphorus and suspended materials in rivers. Water Resources Research 15, 1354–1360 (1979).

14. H. E. Jobson, Predicting travel time and dispersion in rivers and streams. Journal of Hydraulic Engineering 123, 971–978 (1997).

15. B. Zhang et al., Phosphorus fractions and phosphate sorption-release characteristics relevant to the soil composition of water-level-fluctuating zone of Three Gorges Reservoir. Ecological Engineering 40, 153–159 (2012).

16. A. Zhou, H. Tang, D. Wang, Phosphorus adsorption on natural sediments: modeling and effects of pH and sediment composition. Water Res 39, 1245–1254 (2005).

17. J. Z. Zhang, X. L. Huang, Relative importance of solid-phase phosphorus and iron on sorption behavior and sediments. Environmental Science and Technology 41, 2789–2795 (2007).

18. S. Daryanto, L. Wang, P. A. Jacinthe, Meta-analysis of phosphorus loss from no-till soils. J Environ Qual 46, 1028–1037 (2017).

19. D. L. Osmond, A. L. Shober, A. N. Sharpley, E. W. Duncan, D. L. K. Hoag, Increasing the effectiveness and adoption of agricultural phosphorus management strategies to minimize water quality impairment. J Environ Qual 48, 1204–1217 (2019).

20. A. M. Michalak et al., Record-setting algal bloom in Lake Erie caused by agricultural and meteorological trends consistent with expected future conditions. Proceedings of the National Academy of Sciences U S A 110, 6448–6452 (2013).

21. R. P. Stumpf, L. T. Johnson, T. T. Wynne, D. B. Baker, Forecasting annual cyanobacterial bloom biomass to inform management decisions in Lake Erie. Journal of Great Lakes Research 42, 1174–1183 (2016).

22. Recommended phosphorus loading targets for Lake Erie. Final Report of the Annex 4 Objectives and Targets Task Team, May 11, 2015 (2015 https://www.epa.gov/sites/production/files/2015-06/documents/report-recommended-phospho-rus-loading-targets-lake-erie-201505.pdf.).

23. D. Scavia, J. V. DePinto, I. Bertani, A multi-model approach to evaluating target phosphorus loads for Lake Erie. Journal of Great Lakes Research 42, 1139–1150 (2016).

24. T. N. Williamson, E. G. Dobrowolski, A. C. Gellis, T. Sabitov, L. Gorman Sanisaca, Monthly suspended-sediment apportionment for a western Lake Erie agricultural tributary. Journal of Great Lakes Research, (2020).

25. D. B. Green, T. J. Logan, N. E. Smeck, Phosphate adsorption-desorption characteristics of suspended sediments in the Maumee River Basin of Ohio. Journal of Environmental Quality 7, 208–212 (1978).

26. D. L. McCallister, T. J. Logan, Phosphate adsorption-desorption characteristics of soils and bottom sediments in the Maumee River Basin of Ohio. Journal of Environmental Quality 7, 87–92 (1978).

27. H. P. Jarvie et al., Increased soluble phosphorus loads to Lake Erie: unintended consequences of conservation practices? Journal of Environmental Quality 46, 123–132 (2017).

28. D. B. Baker et al., Lagrangian analysis of the transport and processing of agricultural runoff in the lower Maumee River and Maumee Bay. Journal of Great Lakes Research 40, 479–495 (2014).

29. G. Matisoff et al., Internal loading of phosphorus in western Lake Erie. Journal of Great Lakes Research 42, 775–788 (2016).

30. J. D. Chaffin, T. W. Davis, D. J. Smith, M. M. Baer, G. J. Dick, Interactions between nitrogen form, loading rate, and light intensity on Microcystis and Planktothrix growth and microcystin production. Harmful Algae 73, 84–97 (2018).

31. C. A. Stow, Y. Cha, L. T. Johnson, R. Confesor, R. P. Richards, Long-term and seasonal trend decomposition of Maumee River nutrient inputs to western Lake Erie. Environ Sci Technol 49, 3392–3400 (2015).

32. R. P. Richards, D. B. Baker, J. P. Crumrine, Improved water quality in Ohio tributaries to Lake Erie: A consequence of conservation practices. Journal of Soil and Water Conservation 64, 200–211 (2009).

33. D. R. Smith et al., Lake Erie, phosphorus, and microcystin: Is it really the farmer’s fault? Journal of Soil and Water Conservation 73, 48–57 (2018).

34. K. Hayhoe, J. VanDorn, T. Croley, N. Schlegal, D. Wuebbles, Regional climate change projections for Chicago and the U.S. Great Lakes. Journal of Great Lakes Research 36, 7–21 (2010).

35. N. S. Bosch, M. A. Evans, D. Scavia, J. D. Allan, Interacting effects of climate change and agricultural BMPs on nutrient runoff entering Lake Erie. Journal of Great Lakes Research 40, 581–589 (2014).

36. M. M. Kalcic et al., Climate Change and Nutrient Loading in the Western Lake Erie Basin: Warming Can Counteract a Wetter Future. Environ Sci Technol 53, 7543–7550 (2019).

37. K. J. Gibbons, T. B. Bridgeman, Effect of temperature on phosphorus flux from anoxic western Lake Erie sediments. Water Res 182, 116022 (2020).

38. R. M. Kreiling, M. C. Thoms, W. B. Richardson, Beyond the edge: linking agricultural landscapes, stream networks, and best management practices. Journal of Environmental Quality 47, 42–53 (2018).

39. F. Worrall, N. J. K. Howden, T. P. Burt, A method of estimating in-stream residence time of water in rivers. Journal of Hydrology 512, 274–284 (2014).

40. X. X. Lu et al., Sediment loads response to climate change: A preliminary study of eight large Chinese rivers. International Journal of Sediment Research 28, 1–14 (2013).

41. R. H. Meade, J. A. Moody, Causes for the decline of suspended-sediment discharge in the Mississippi River system, 1940-2007. Hydrological Processes, n/a-n/a (2009).

42. S. V. Mize, J. C. Murphy, T. H. Diehl, D. K. Demcheck, Suspended-sediment concentrations and loads in the lower Mississippi and Atchafalaya rivers decreased by half between 1980 and 2015. Journal of Hydrology 564, 1–11 (2018).

43. B. J. Peterson et al., Controls of nitrogen export from watersheds by headwater streams. Science 292, 86–90 (2001).

44. P. A. Raymond et al., Global carbon dioxide emissions from inland waters. Nature 503, 355–359 (2013).

45. H. P. Jarvie et al., Within-river phosphorus retention: accounting for a missing piece in the watershed phosphorus puzzle. Environ Sci Technol 46, 13284–13292 (2012).

46. M. J. White et al., Development and testing of an in-stream phosphorus cycling model for the Soil and Water Assessment Tool. Journal of Environmental Quality 43, 215–223 (2014).

47. H. P. Jarvie et al., Quantifying phosphorus retention and release in rivers and watersheds using extended end-member mixing analysis (E-EMMA). Journal of Environmental Quality 40, 492–504 (2011).

48. H. W. Paerl et al., It Takes Two to Tango: When and Where Dual Nutrient (N & P) Reductions Are Needed to Protect Lakes and Downstream Ecosystems. Environmental Science & Technology 50, 10805–10813 (2016).

49. J. C. Finlay, G. E. Small, R. W. Sterner, Human influences on nitrogen removal in lakes. Science 342, 247–250 (2013).

50. K. L. Blann, J. L. Anderson, G. R. Sands, B. Vondracek, Effects of agricultural drainage on aquatic ecosystems: a review. Critical Reviews in Environmental Science and Technology 39, 909–1001 (2009).

51. Ohio EPA, Nutrient mass balance study for Ohio’s major rivers. Ohio Environmental Protection Agency Division of Surface Water, Ed., (Ohio Environmental Protection Agency Division of Surface Water,, Columbus, OH, 2018).

52. J. D. H. Strickland, T. R. Parsons, A practical handbook of seawater analysis. (Fisheries Research Board of Canada, Ottawa, 1972).

53. H. P. Jarvie et al., Role of river bed sediments as sources and sinks of phosphorus across two major eutrophic UK river basins: the Hampshire Avon and Herefordshire Wye. Journal of Hydrology 304, 51–74 (2005).

54. D. Y. F. Lai, K. C. Lam, Phosphorus sorption by sediments in a subtropical constructed wetland receiving stormwater runoff. Ecological Engineering 35, 735–743 (2009).

55. P. N. Froelich, Kinetic control of dissolved phosphate in natural rivers and estuaries: A primer on the phosphate buffer mechanism. Limnology and Oceanography 33, 649–668 (1988).

56. U.S. Geological Survey [USGS]. (2021), vol. 2021.

57. G. Fouad, A. Skupin, C. L. Tague, Regional regression models of percentile flows for the contiguous US: Expert versus data-driven independent variable selection. Hydrology and Earth System Sciences, (2016).

58. H. U. National Center for Water Quality Research. (2020).

59. S. N. Wood, Generalized additive models: an introduction with R. (Taylor & Francis, Boca Raton, FL USA, 2006).

60. S. N. Wood. (R Core Team 2017, 2019).

61. R Core Team. (R Foundation for Statistical Computing, Vienna, Austria, 2020).

62. J. Du, H. Xie, Y. Hu, Y. Xu, C.-y. Xu, Development and testing of a new storm runoff routing approach based on time variant spatially distributed travel time method. Journal of Hydrology 369, 44–54 (2009).

63. H. H. Barnes, Jr.,, Roughness characteristics of natural channels. U. S. G. Survey, Ed., (U.S. Geological Survey, Denver, CO, 1967).

64. K. P. Burnham, D. R. Anderson, Model selection and multimodel inference: A practical information-theoretic approach. (Springer, New York, ed. 2nd Edition, 2002).

65. D. I. Warton, R. A. Duursma, D. S. Falster, S. Taskinen, smatr 3 - an R package for estimate and inference about allometric lines. Methods in Ecology and Evolution 3, 257–259 (2012).

66. R. L. Wasserstein, A. L. Schirm, N. A. Lazar, Moving to a World Beyond “p < 0.05”. The American Statistician 73, 1–19 (2019).

